# Electrode sharpness and insertion speed reduce tissue damage near high-density penetrating arrays

**DOI:** 10.1101/2023.11.22.568119

**Authors:** Ingrid N. McNamara, Steven M. Wellman, Lehong Li, James R. Eles, Sajishnu Savya, Harbaljit S. Sohal, Matthew R. Angle, Takashi D. Y. Kozai

## Abstract

**Objective:** Over the past decade, neural electrodes have played a crucial role in bridging biological tissues with electronic and robotic devices. This study focuses on evaluating the optimal tip profile and insertion speed for effectively implanting Paradromics’ high-density Fine Microwire Arrays (FμA) prototypes into the primary visual cortex (V1) of mice and rats, addressing the challenges associated with the “bed-of-nails” effect and tissue dimpling.

**Approach:** Tissue response was assessed by investigating the impact of electrodes on the blood-brain barrier (BBB) and cellular damage, with a specific emphasis on tailored insertion strategies to minimize tissue disruption during electrode implantation.

**Main Results:** Electro-sharpened arrays demonstrated a marked reduction in cellular damage within 50 μm of the electrode tip compared to blunt and angled arrays. Histological analysis revealed that slow insertion speeds led to greater BBB compromise than fast and pneumatic methods. Successful single-unit recordings validated the efficacy of the optimized electro-sharpened arrays in capturing neural activity.

**Significance:** These findings underscore the critical role of tailored insertion strategies in minimizing tissue damage during electrode implantation, highlighting the suitability of electro-sharpened arrays for long-term implant applications. This research contributes to a deeper understanding of the complexities associated with high-channel-count microelectrode array implantation, emphasizing the importance of meticulous assessment and optimization of key parameters for effective integration and minimal tissue disruption. By elucidating the interplay between insertion parameters and tissue response, our study lays a strong foundation for the development of advanced implantable devices with a reduction in reactive gliosis and improved performance in neural recording applications.

## Introduction

Intracortical electrodes are an essential front-end tool for brain-computer interfaces (BCIs)^1,2^. These technologies have enabled a deeper understanding of brain function through fundamental neuroscience research^3-6^ and showcasing promising potential for the restoration of functional motor control in individuals with tetraplegia as well as sensory restoration^7-20^.

Penetrating intracortical electrodes have the potential to offer finer spatial and temporal resolution neural data crucial for high degrees of freedom applications, surpassing the capabilities of surface (ECoG) or endovascular microelectrodes^21,22^. Despite their promising capabilities, the long-term efficacy of these neural electrodes is often compromised by tradeoffs in challenges related to their integration into host tissue^23-30^, limited channel count^31^, low recording site densities^31,32^, and small recording radiuses^2,33^. These limitations have prompted researchers to explore innovative solutions to enhance the performance of these devices, paving the way for advancements in the development of high-density electrode arrays^34^.

Early studies of cortical reactive gliosis indicate that the size of implantable devices did not impact the overall foreign body response^35^. However, recent studies have emphasized the significance of “subcellular-sized” microelectrodes in mitigating the adverse tissue responses typically associated with conventional-sized electrodes^36^. This reduced tissue response is attributed to the diminutive dimensions of these microelectrodes, which facilitate improved neural density around the probes, a decrease in gliosis and scar tissue formation, and improved recording performance^37,38^. In turn, this research has underscored the critical role of design considerations, particularly due to the steep signal drop-off^2^ (1/r∼1/r^2^) of recorded actions potentials that occurs with microelectrodes that typically have a small recording radius of ∼100 µm^33^ even with low impedances^38^. As expected, functional microelectrodes with subcellular dimension made from silicon or carbon demonstrated improved recording performance compared to traditionally sized devices^38-43^. However, a number of technical hurdles remain for implementing these technologies on a high-channel count scale^43^.

The implementation of high-channel-count arrays on a large scale poses substantial technical challenges^41-43^. One of the primary hurdles is the manual assembly process associated with ultra-small carbon fiber devices, making it impractical to build arrays with hundreds or thousands of channels efficiently^44^. Furthermore, the transition to subcellular-sized devices has presented an additional trade-off, with a need to address the increased fragility of these devices^2,45^. Strategies involving cylindrical geometries devoid of hard corners have been proposed to mitigate this issue^2,46^. However, blunt probes pose several challenges during tissue penetration, including increased insertion forces, a higher likelihood of missing anatomical targets, and heightened potential for tissue trauma^47-50^. Additionally, the crucial phase of electrode insertion demands careful consideration, as sharper tip profiles and controlled insertion speeds have been shown to be instrumental in preserving tissue integrity and optimizing electrode performance^51-53^. Although 16-channel Utah arrays can easily be inserted at a slow speed^32^, 100-channel arrays need to be inserted at ballistic speeds using a pneumatic inserter ^54^. With an increasing number of channels, it becomes increasingly more difficult to insert bed-of-needle arrays (microscale “bed-of-nails”) without compressing and damaging the underlying brain tissue.

Once penetration through the brain surface is achieved, implantation of intracortical arrays leads to severing and rupturing of blood vessels ^55-57^; tearing of neuronal cell membranes and calcium activation^58^; injury to axons and myelin ^29,59,60^; and activation of glial cells such as microglia, astrocytes, and oligodendrocyte progenitor cells^24-26,61-68^. In addition, as the number of channels increases, the total volume of the device that is implanted also increases, requiring the displacement of a greater volume of brain tissue^2^. As the pitch of individual shafts is made smaller, the tissue response eventually treats the individual sub-cellular shafts as one large electrode shaft^43^. Careful consideration of interdependent design parameters, together with innovative design strategies, are necessary for the practical and functional implementation of high-density, high-channel-count arrays^2^.

Furthermore, the challenges associated with the practical implementation of high-channel-count arrays extend to the intricate aspects of device connectivity and the effective extraction of neural signals amidst external noise sources. The limitations imposed by connector bandwidths and computational power necessitate innovative approaches, including the integration of onboard multiplexers to streamline data transmission^34^. However, the use of multiplexers introduces capacitively coupled artifacts in the recording data, constraining the sampling frequency of the channels and posing limitations on the comprehensive recording of wideband data. To address these concerns, Paradromics, Inc. has embarked on developing a scalable, high-throughput strategy for multichannel electrophysiology, emphasizing the critical role of a viable front-end interface that seamlessly integrates with the brain tissue while upholding the functionality of nearby neurons^69^.

Nevertheless, the success of this strategy is predicated on the success of a front-end interface successfully implanting into the brain while maintaining viability of nearby neurons. The current study seeks to evaluate the efficacy of Paradromics’ front-end prototype arrays (Fine Microwire Arrays, FμA) through an extensive, iterative assessment of various parameter spaces, aimed at achieving a high-channel-count microelectrode array. Particularly, we focus on bypassing the “bed-of-nails” effect, where the insertion force of the electrode array is distributed over a larger surface area, diminishing the ability for any single electrode in the array to penetrate the tissue and leading to dimpling of the brain. The evaluation process entails a comprehensive examination of multiple pitch sizes, tip profiles, and insertion speeds conducive to successful intracortical insertion. A central objective of this study is to meticulously characterize the tissue response to the microelectrode array, facilitating a high-throughput evaluation across diverse parameter configurations. This characterization involves the assessment of cell membrane rupture through the use of propidium iodide (PI) and the analysis of blood-brain barrier (BBB) integrity and leakage via immunoglobulin (IgG) staining. Moreover, a comparative evaluation between the FμA and commercially available Blackrock arrays is conducted, shedding light on the specific array configurations and insertion parameters that minimize the impact on BBB and surrounding tissue from the electrode sites.

## Materials and Methods

### Microelectrodes

Fine microwire electrode prototype arrays (FμA) (∼24 channels or ≥60 channels) with a diameter of 20 μm for each electrode was provided by Paradromics, Inc., Austin, TX, USA (paradromics.com) and tested to address optimal insertion parameters for larger bundle arrays. Proprietary prototype arrays were microwires with glass insulation as previously described^70^. FμA were provided with three different tip profiles: blunt, angled cut (30-degree angle), and electro-sharpened with a short pinpoint tip (∼8-degree angle) (Figure 1, A)^70^. The blunt and angle Au arrays had a 100 μm pitch (∼100 electrodes); electro-sharpen W arrays had 300 μm pitch (∼160 electrodes); and control (16 electrodes) with 400 μm pitch. These FμA were tested (**Fig. 1A-D**) and compared to Utah Electrode Array (UEA) (Blackrock Microsystems, Salt Lake City, UT) as the control array (**Fig. 1E**). W was chose to replace Au wires for electrosharpened arrays, due to concerns that Au would be too soft to maintain the sharpened shape. In addition, to test the recordability of the individual wires of the FμA implants, single wire electrodes (20 μm in diameter) were implanted and electrophysiologically evaluated.

**Figure 1:**
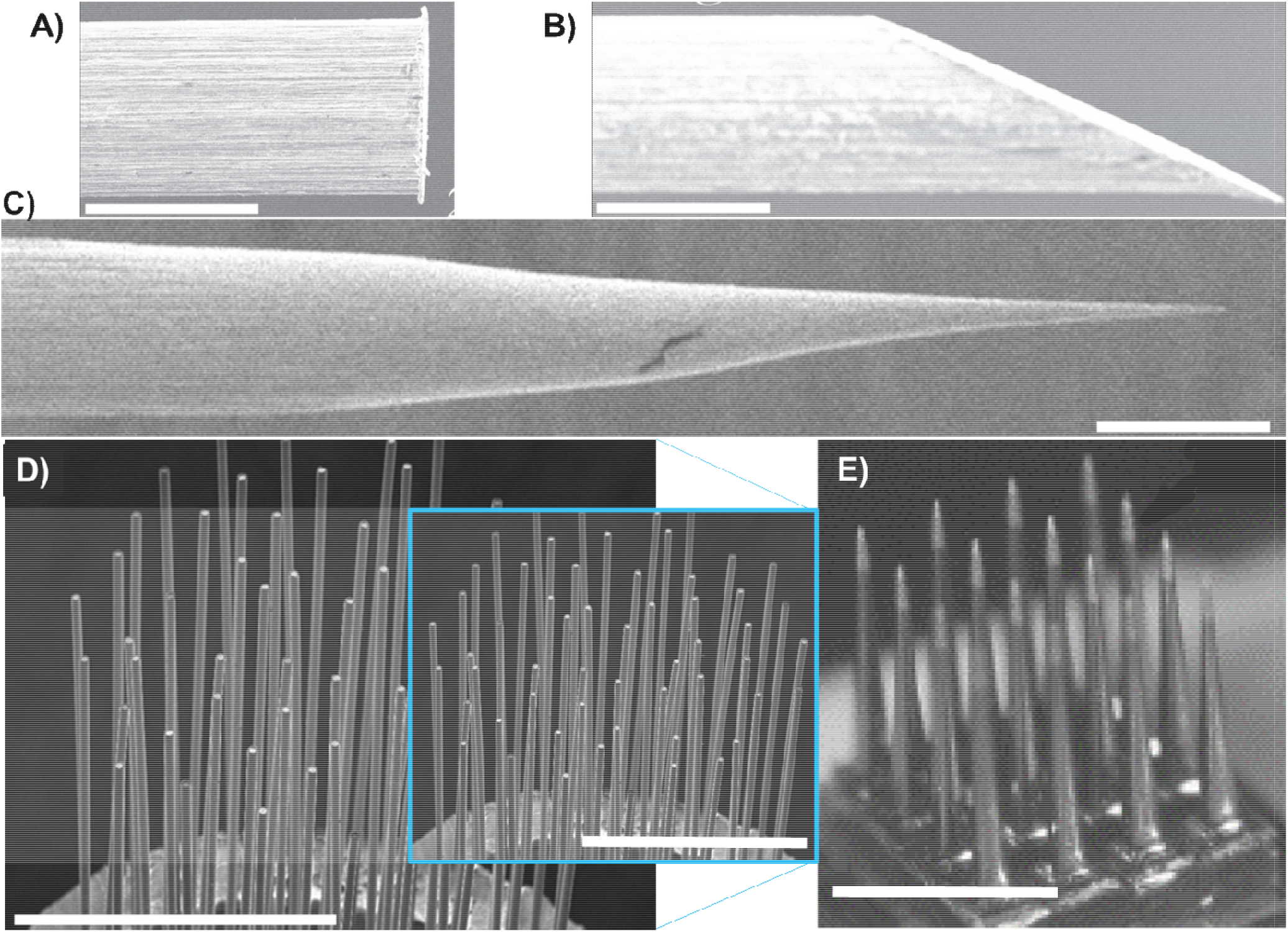
Microelectrodes. **A-C)** Schematic of the tip profiles: blunt (**A**), angle (∼30 degree cut) polished (**B**), and electro-sharpen (**C**). Scalebar = 25 µm. **D)** Image shows an example of a high-density array (1.6 mm in diameter) with about ∼55 individual electrodes with 18 μm (+/-0.5) diameter gold glass wires with 100 μm electrode spacing (pitch) and 1.5 m in length. Scalebar = 1 mm. Cyan box shows the same array to scale to the Blackrock array (**E**). **E)** Image of 4×4 Blackrock array with 400 μm spacing and tapered 25 μm diameter electrodes (1.2 mm x 1.2 mm). Scalebar =1 mm .

### Surgical Procedures for probe insertion in mice for two-photon imaging

Seven mice were induced with 75ml/kg of Ketamine with 7 mg/kg Xylazine via an intraperitoneal (IP) injection and the skull was then shaved and prepped for surgery. The animal was maintained with ketamine (40 mg/kg) so that it did not respond to a toe pinch and remained properly anesthetized throughout the experiment. The animal was placed in the stereotaxic frame and on top of a heating pad with a rectal probe for maintaining the proper body temperature (37^°^C). An ocular ointment was placed on the eyes to prevent desiccation. Next, the area was prepped with three cycles of the betadine scrub and alcohol washes. The skin was removed over the skull and the connective tissue was reflected back and bone exposed. Vetbond (3M) was placed over the bone surface and two bone screws were placed over the frontal cortex areas. Dental cement was used to anchor the bone screws and to create an imaging well for holding the saline for the water-immersion objective lenses. Next, a 4 mm by 6 mm craniotomy was made over the visual cortex centrally located over 1.5 mm lateral from lambda and 1.5 mm rostral. The implant area was frequently flushed with sterile saline to prevent thermal heating due to drilling and prevent the tissue from desiccating during the procedure.

FμA probes (∼24 channel FμA) were inserted at a 30-35 degree angle from the horizontal plane and parallel to midline down to layers II/III (∼300 μm down from the pia) in the mice visual cortex area. The arrays were inserted with a Narishige oil hydraulic drive (MO-81, Narashige, Japan) at a speed of 500 nm s-1. For animals that were imaged for cellular damage, propidium iodide (PI, 1mg/mL in saline) was placed on the surface of the brain post insertion. After 20 minutes the surface of the brain was flushed 3 times with saline prior to imaging. Also, for the electro-sharpen insertions sulfonamide (SR101) was injected (IP) for visualizing the vasculature around the electrode sites. Images were acquired at 1024 × 1024 pixel (407.5 × 407.5 μm) over ∼4.8 s using Prairie View software. Stacks were acquired along depth (ZT stacks) every 2 μm from the surface to 100 microns below the tip of the electrode. Images were taken from the individual Z stacks to calculate the number of damaged cells around the implanted electrode wires and were measured using the ‘Measure’ function in ImageJ (National Institutes of Health). All procedures were approved by the Division of Laboratory Animal Resources and the Institutional Animal Care and Use Committee of the University of Pittsburgh and in accordance with the standards set by the Animal Welfare Act and the National Institutes of Health Guide for the Care and Use of Laboratory Animals.

### Two-photon imaging

Two-photon microscopy techniques were used to examine the cellular responses to tip profile under *in vivo* conditions in mice and followed the similar procedures in previous work^64,66-68,71-77^. The microscope for the *in vivo* imaging used a scan head (Ultima I.V.; Bruker, Madison, WI), an OPO laser (Insight DS+; Spectra-Physics, Menlo Park, CA) tuned to a wavelength of 920 nm, non-descanned photomultiplier tubes (Hamamatsu Photonics KK, Hamamatsu, Shizuoka, Japan), and a 16X, 0.8 numerical aperture water immersion objective lens (Nikon Instruments, Melville, NY). Transgenic mice (25-35g, Jackson Laboratories) were used that expressed green fluorescent protein (GFP) in the microglia (B6.129P-cx3cr1/J, n=3) or in neurons (Thy1-GCaMP3)4.3 for imaging (n=3). In addition, propidium iodide (PI, 1 mg/mL in saline) was used for identifying damaged cells and sulfonamide (SR101, 0.02–0.04[cc; 1[mg of drug per ml of sterile saline) was used to localize blood vessels^78^.

### Surgical Procedures in rats for Perpendicular Electrode Implantation

Acute Experiments were performed on 19 Sprague Dawley (Charles, River), male rats (400+50 g) and implanted with non-functional high-density arrays (≥60 electrodes) in each hemisphere as previously described^59^. For 1-week chronic experiments, 8 animals were implanted. Prior to surgery animals were given 75 mg/kg Ketamine and 7 mg/kg Xylazine cocktail via an intraperitoneal (IP) injection with regular updates of 40 mg/kg Ketamine IP. The anesthetic level was confirmed via the absence of a reaction from toe pinch response. The surgical site was shaved and the animal was placed in the stereotaxic frame (David Kopf Instruments, CA) and vitals were monitored throughout the procedure. Proper body temperature was maintained (37 □ C) via a heating pad. Next, the ocular ointment was placed on the eyes and the surgical area was prepped with betadine surgical scrub and then cleaned with 70% isopropyl alcohol. An incision was made over the scalp exposing the surgical area. The fascia was reflected and the periosteum from the bone was removed and a thin layer of cyanoacrylate (Vetbond, 3M) was placed over the skull. A pinhole craniotomy was made with a manual drill and rongeurs were used to perform the craniotomy. The tissue was frequently irrigated to prevent desiccation and wash away bone debris. The craniotomy was placed over the V1 area approximately -5mm to -7mm of bregma and -2mm to -4mm lateral from midline and about 5mm x 5mm in dimensions. After the durotomy, a micromanipulator was used to insert the electrodes into the brain and physiological saline was used to keep the brain moist during the procedures. The arrays were inserted at a 90^°^ angle into the brain approximately 1mm down with either the slow, fast or pneumatic insertion method (as described in the following section). After the electrodes were inserted, propidium iodide (PI, 1 mg/mL in saline) was placed over the surface of the cortex for 20 minutes to label damaged or compromised cell membranes (n=15 of acutely implanted animals), a method used in previous studies. Following the incubation period, the PI ^60,79^ was rinsed with three flushes of saline; the craniotomy was sealed with dental cement (Henry Schein, Flowable Composite 101-6773). Then, the animals were transcardially perfused with PBS and then with 4% paraformaldehyde for fixation ^79-82^.

### Insertion Methods in rats

Fast insertions were performed using a linear translator by Physik Instrumente (V551-4B stage from Physik Instrumente). Probes were positioned at -22.162 mm from the brain surface to achieved a velocity of 80 mm/sec at the time of contact with the brain surface, which was followed by a deceleration length of < 0.75mm (Supplementary Figure 1). Slow insertions were performed with a manual micromanipulator and inserted slowly by hand at approximately 1-2 mm/s.

Also, the pneumatic inserter (Blackrock Microsystems) was used to compare tissue response to various insertion methods. The pneumatic insertion uses a vacuum pump to reach very high velocities for inserting surface probe arrays ^32,82^. Proper calibration of the pneumatic inserter was done to only allow a “1 hit” insertion, preventing a “double-hit”. All electrodes were zeroed to the surface on the pia and then advanced 1mm down into the brain. The control array was tested across slow and pneumatic insertions and used as a comparison to the other tested methods ^32^.

### Histological Processing

After the animal was perfused, the skull was dissected and placed in a 4% paraformaldehyde for up to ∼12 hours. Next, the brain was dissected out and placed in 15% sucrose until tissue dropped (∼12-18 hours). Next, the brain was transferred to 30 % sucrose solution until the tissue reached equilibrium. Once the tissue dropped in the 30% sucrose (∼1-2 days), the tissue was then blocked and frozen in a mold in Optimal Cutting Temperature (OCT) compound 2:1 mixture (20% sucrose: OCT) on a shallow dish of 2-methylbutane sitting on dry ice. Finally, the tissue was sectioned serially (15μm thick horizontal sections) and stored in -20^°^C until staining.

### Immunostaining & Fluorescence Imaging

Sections were re-hydrated within a week with 1XPBS for 15 minutes followed by the following stains: Hoechst 33342, Nissl (500/525) and Donkey anti-rat Immunoglobulin G (IgG) for analysis. Slides were coverslipped with Fluoromount-G and stored at 4^°^C in the dark until imaged within the following week. Images were acquired with a Leica microscope and processed using ImageJ for analysis with the I.N.T.E.N.S.I.T.Y. Analyzer ^25,83^. The sections were analyzed with the radial image binning around the probe site to calculate the changes in fluorescence intensity from the IgG. The background intensity for each image was calculated by the 4 corners within each image. For all pixels with less than 1 standard deviation (STD) of the mean intensity was used as the background threshold and the image threshold was set to 1 STD. The Region of Interest (ROI) was selected and binned at 10 μm with a total of 32 bins and the electrode diameter was 20 μm for each image. For the control array, the electrode diameter was set to 25 μm and binned at 10 μm with a total of 32 bins. The single wire electrodes that were inserted were 20 μm and the control (Microprobe single wire) was 81 μm in diameter. The single wires were also binned at 10 μm with 32 bins for image analysis.

### Electrophysiology

Sprague-Dawley (SD) rats (n=3) were induced with Ketamine and Xylazine and maintained with Ketamine as listed under surgical procedures. The electrophysiological recordings were taken inside a Faraday cage while single wire electrodes were implanted into the left visual cortex and a monitor was placed outside the cage in the right visual field as described in previous studies^3,28,30,32,39,44,84-88^. The electrode impedance for the electrode was 0.17 Mohm at 1 kHz and recordings were made at multiple depths (0, 520 and 1000 μm) down in the brain. Tucker Davis Technologies system (Medusa preamp, Tucker-Davis Technologies, Alachua FL) was used to record the cortical signals (high pass 300 Hz and low pass at 5000 Hz) for 3-minute trials. The visual stimulus was made up of 8 different directional translations of drifting gratings that moved across the monitor (1 sec) while the neural response was recorded from a single wire (20 μm in diameter) electrode^85^. The 8 directional drifting grating stimulation were repeated 8 times, for a total of 64 stimulation trials. Once units were sorted as previously described^37^, the firing rate was determined for each unit and unsorted multiunit threshold crossings. The units were sorted into 50 ms binned PSTH from -1 s before the visual stimulation to 1 s after the stimulation. The visual cortex was chosen for assessment, allowing the evaluation of functional connectivity of recorded single-units through visual stimulation to the contralateral retina under acute anesthetized conditions. This choice ensures confidence in distinguishing between functional evoked neural activity and potential spontaneous dysfunction or epileptic neural activity due to ruptured axons from insertion strain that potential disconnected neurons near the electrode from the broader neural network. This distinction would not be possible in M1 under acute anesthetized condition.

### Data Analysis

Intensity values were extracted from I.N.T.E.N.S.I.T.Y. Analyzer per image (≥ 2 animals and ≥ 3 electrode sites per condition) and reported as a mean ± standard error as a function of distance from the insertion probe site. The intensity data was averaged over 150 μm away from the electrode site and generated bar graphs reporting the mean ± standard error for each insertion condition. Also, the number of damaged cells were counted manually by identifying neural and cell body staining (Hoechst/Nissl) that showed co-localization staining with PI. The ImageJ measurement tool was used to count and calculate the distance for each damaged neuron from the center of the electrode site. Measurements were taken 0 to 150 μm away from the center of the insertion site for the Two-Photon *in-vivo* images. Cell count images were binned 50 μm up to 150 μm away from the center of the electrode site. Cell counts were averaged per bin and reported as a mean ± standard error as a function of distance from the electrode site.

### Statistics

Bar plots were compared using an unequal variance Welch’s t-test followed by a Bonferroni correction to correct for repeated measures. In addition, a two-way ANOVA with a post-hoc Tukey test was performed on line plots to examine interactions between the insertion parameters: tip profiles (blunt, angle, electro-sharpen, and control) and speeds (slow, fast, and pneumatic).

## Results

The study focused on assessing insertion strategies for implanting high-channel-count microelectrode arrays (FμA) with specific dimensions (≥60 channels, ∼20 μm diameter, ∼100 to 300 μm pitch), employing three distinct tip profiles and three insertion speeds. Successful insertions were determined based on the identification of probe tracks in cortical layers housing neuronal cell bodies. Additionally, the investigation included the evaluation of cell membrane rupture and blood-brain barrier (BBB) leakage to quantify acute insertion-related injuries, facilitating the identification of optimal array and insertion parameters. Furthermore, the study aimed to verify the suitability of the final electrode design parameters for the array by conducting single-unit recordings from individual FμA wires, ensuring their compatibility with electrophysiological recording requirements.

### In vivo two-photon insertion in mice

Implantation of microelectrodes is known to induce tissue compression^45^ and compromise neuronal membranes, as evidenced by membrane-impermeable dyes such as PI^57^. Given that multiple shanks of an array can contribute to overall tissue strain^2^ in a geometrically dependent manner^45^, we sought to understand how the tip geometry of high-density arrays influences acute neuronal injury. To address this, we compared various tip profiles, including blunt, angle polished (30-degree angle cut), and electro-sharpened ∼24 channel arrays during in vivo two-photon insertion analysis. *In vivo* images highlighted damaged cells (PI+) and Ca^2+^ active neurons around the electrode tip profiles (**Fig. 2A**). The findings indicated a lower incidence of membrane injured cells around the electro-sharpened tips, while the blunt and angled tips caused significantly greater membrane damage around the insertion site (unequal variance t-test *p*<0.05; **Fig. 2B**). Taken together, implanting electro-sharpened tipped arrays *in vivo* exhibited less cellular injury as indicated by PI labeling in the first 50 μm from the electrode tip compared to blunt or angle polished tip profiles (**Fig. 2B**).

**Figure 2:**
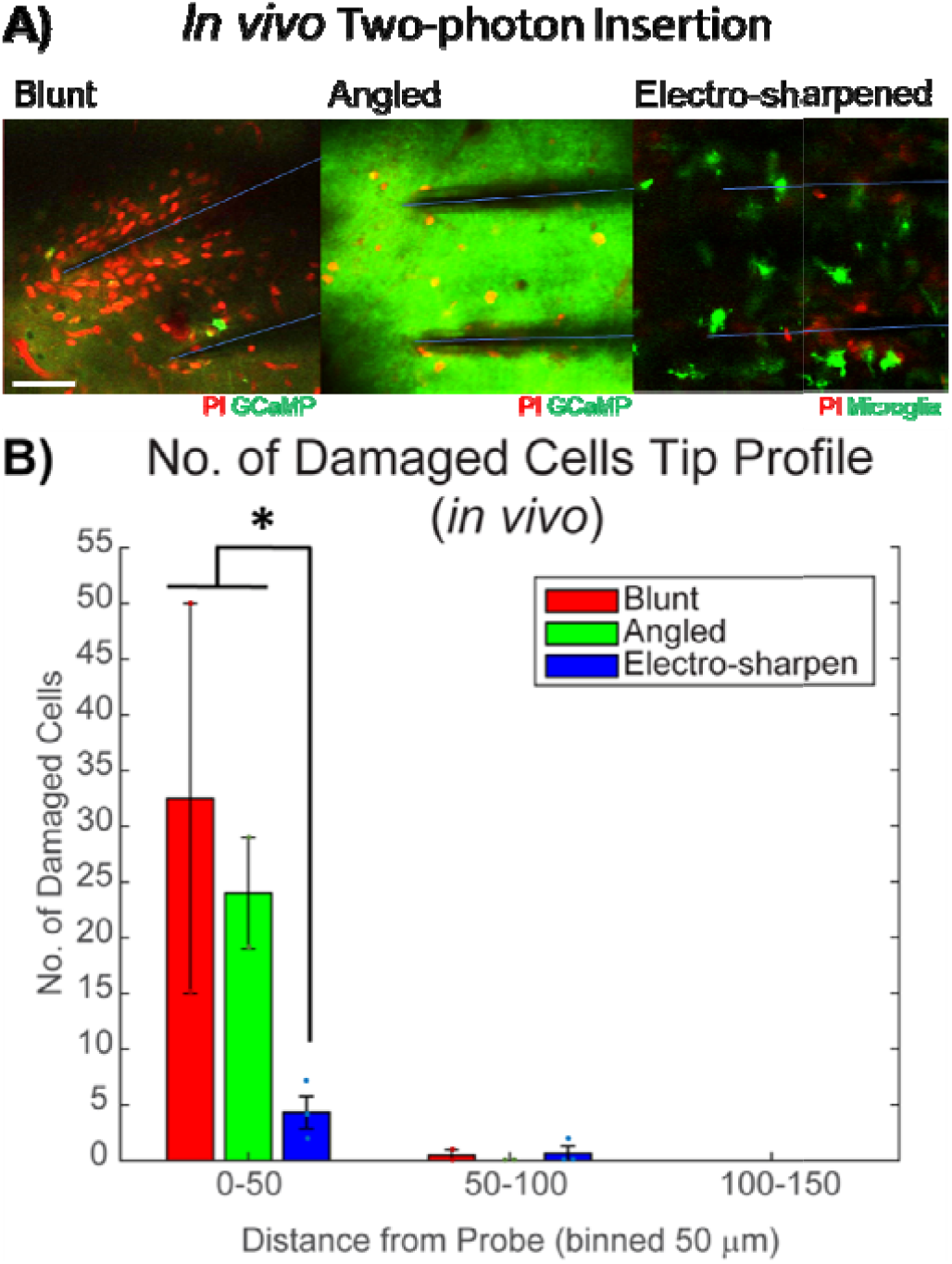
**A)** Two-photon *in vivo* microscopy was performed for different tip profiles (blunt, angle and electro sharpen). Green: Neurons (blunt/angle); Microglia (electro sharpen). Red: cells stained with PI indicating damaged ells; and sulfonamide (SR101) was used to localize blood vessels. Blue dotted line indicates the location of the electrode shanks. Electrodes were inserted at ∼30 degree an le so that the electrode could be viewed during insertion. The results show that there are less injured cells around the electro-sharpen arrays compared to the angle and blunt ti profiles. The white scale bar = 100 μm. **B)** The bar plot hows the distribution of damaged cells from the electrode shank. The blunt and angled tip profile indicates the greatest mount of PI stained cells. The electro-sharpen insertion had the least amount of damage to the cells around the impla t sites. ^*^ indicates p<0.05 with an unequal variance t-test. (: Blunt = 2, Angled = 2, electro-sharpened = 3).

### Tip profile -Angle, Electro-sharpened, and Control in rats

Given the efficacy of electro-sharpened arrays in reducing neuronal injury during angled insertion in mice, we aimed to determine whether these results held true for larger channel arrays implanted perpendicularly into rats. To address this, we evaluated the same electrode tip profiles integrated into larger arrays (≥60 channels) by measuring the disruption of the blood-brain barrier (BBB) leakage with IgG and cellular damage during insertion of neurons with Nissl+PI (**Fig. 3A**). All three profiles and the control Blackrock arrays were inserted at the same slow speed (1-2 mm/s) to ensure an unbiased analysis of the electrode profiles. Neurons were labeled with Nissl, while PI was used to identify cells with compromised cellular membranes. The co-localization of Nissl and PI staining highlighted the areas of damaged cells or neurons, indicated by the yellow arrow (see **Fig. 3A**). Notably, the blunt tips did not insert and only caused dimpling of the tissue. The other large bundles were successfully inserted, although all arrays showed some tissue dimpling in the histological images. Careful attention was paid to avoid post-processing artifacts and prevent any tissue damage during array removal. The analysis indicated minimal IgG differences among the three profiles within 0 – 150 μm from the electrode site (p>0.05; **Fig. 3B**), suggesting a comparable level of BBB injury. However, it should be noted that a substantial outlier in IgG leakage was induced by one of the large Blackrock shanks, attributed to its penetration of an invisible arteriole.

**Figure 3:**
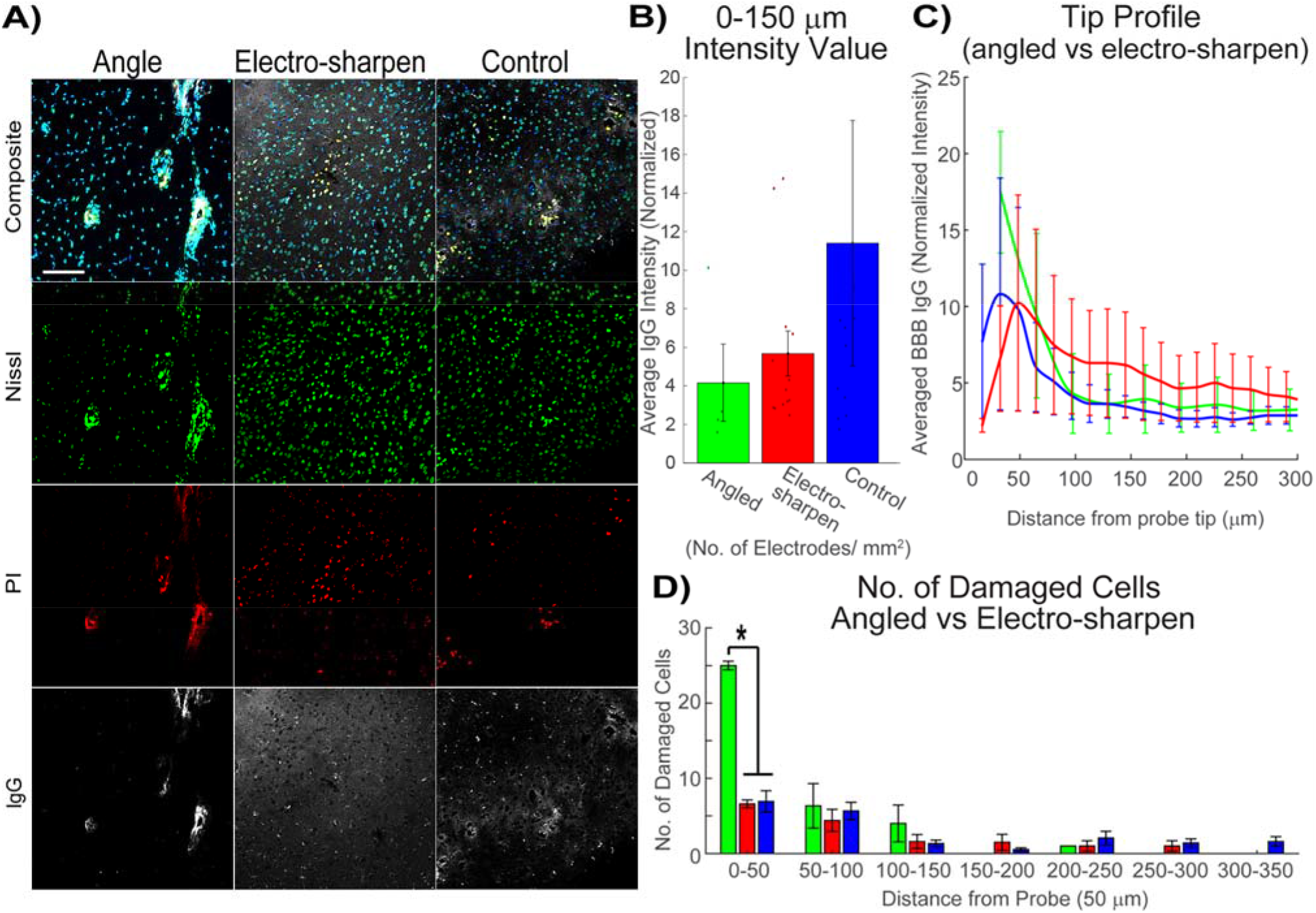
**A)** Fluorescence staining for Hoechst stain (Blue); Nissl (Green); PI (Red); and IgG (White) shows the tissue response to the different electrode tip profiles (angle, electro-sharpen and control). The control array (Blackrock array) and all tip profiles were inserted at the same slow speed. The yellow arrows identify example area of co-localization for the Nissl and propidium iodide (PI) stained cell and indicated areas of damaged cells or neurons. White scale bar = 100 μm. **B)** Normalized IgG intensity within 0 – 150 μm from probe hole. **C)** Normalized IgG intensity within 10 μm bins up to 300 μm away from the probe hole. **D)** The number of damaged cells (PI+) labeled was measured up to 350 μm away from the probe hole and averaged within 50 μm bins. ^*^ indicates p<0.05 with an unequal variance t-test. (N: angled = 4, electro-sharpened = 5, control = 2. N_sites_: angled = 4, electro-sharpened = 13, control = 11)

Furthermore, the analysis did not reveal a significant difference between the electrode profiles in terms of the BBB response (p>0.05; **Fig. 3C**). However, the angled tip exhibited a significantly higher number of PI+ labeled cells compared to the other tip profiles within the first 50 μm of the electrode site (p <0.05). Nonetheless, no significant differences were observed between the bins and groups (p>0.05; **Fig. 3D**). Taken together, the results highlight the role of tip profiles and insertion dynamics in influencing the number of damaged cells during electrode implantation, demonstrating the potential benefits of tailored insertion strategies in minimizing acute tissue damage and enhancing the efficacy of high-density array integration.

### Insertion speed – Slow vs Fast vs Pneumatic in Rats

Given that tip profiles of high-density arrays did influence dimpling, insertion, and neuronal membrane injury, we further investigated the impact of insertion speed on BBB injury and neuronal membrane damage. Prior research focused mainly on single shank microelectrodes, making it important to examine this relationship in the context of array insertions^45,51^.

Three insertion speeds, namely slow (1-2 mm/s), fast (80 mm/s), and pneumatic (∼200 mm/sec), were evaluated for their effects on BBB leakage (IgG) and neuronal membrane damage (Nissl and PI) (**Fig. 4A**). Co-localization of Nissl and PI staining revealed the regions of cellular damage or neuronal impairment (**Fig. 4A**). The angled tip profile arrays (30-degree bevel cut tip) were used for comparing the various insertion speed as these arrays were more abundantly available due to the ease of fabrication compared to the proprietary electro-sharpened arrays. Slow insertion speed demonstrated the greatest amount of BBB leakage compared to the fast and pneumatic methods. In contrast, IgG intensity values within 0 – 150 μm from the electrode sites showed the lowest values with the pneumatic insertion, which demonstrated notably higher insertion velocity. While the study did not find a significant difference between slow and fast insertion methods (p>0.5), slower speeds were found to significantly impact the BBB compared to pneumatic insertions (p<0.05; **Fig. 4B**). Moreover, the IgG intensity levels varied significantly among the groups (p<0.05) but not between the bins and groups (p>0.05; **Fig. 4C**). Interestingly, the pneumatic insertion method elicited a comparatively lower BBB response, but the area surrounding the implant site exhibited a greater number of PI-stained cells (p<0.05; **Fig. 4D**). Taken together, our findings underscore the complex interplay between insertion dynamics, tip profiles, and insertion speeds, emphasizing the necessity for meticulous consideration of these parameters to minimize tissue damage.

**Figure 4:**
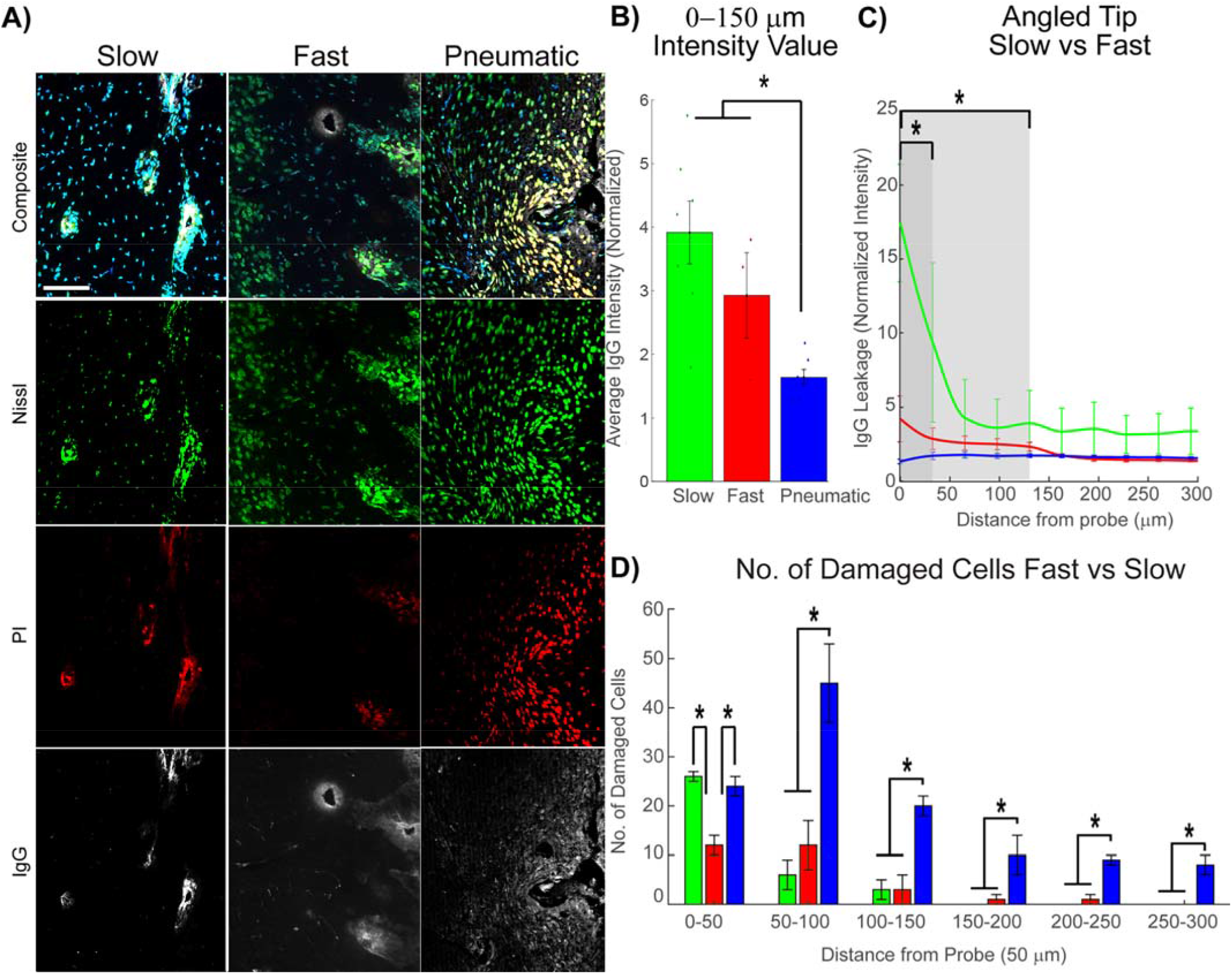
**A)** Fluorescence staining for Hoechst stain (Blue); Nissl (Green); PI (Red); and IgG (White) displays the tissue response to the different insertion speeds (slow, fast and pneumatic). The control speed was the pneumatic insertion method and all array profiles were the angle tip profiles within the different insertion speeds. The yellow arrows identify example area of co-localization for the Nissl and PI stained cell and indicated areas of damaged cells or neurons around the insertion site. White scale bar = 100 μm. **B)** Normalized IgG intensity within 0 – 150 μm from probe hole for the different insertion speeds. (N_electrode_: Slow= 7, Fast = 3, Pneumatic = 6). **C)** Normalized IgG intensity within 10 μm bins up to 300 μm away from the probe hole. **D)** The number of damaged cells (PI+) measured up to 300 μm away from the probe hole and averaged within 50 μm bins. ^*^ indicates p<0.05 with an unequal variance t-test. (N_rats_: Slow = 4, Fast = 3, Pneumatic = 2)

### Chronic Response to Large Bundle Insertions -Slow vs Fast & Acute Response in Rats

Given that slow and fast insertion led to the least amount of neuronal membrane injury, we proceeded to investigate whether electro-sharpened arrays insertion speeds led to increased BBB leakage one week post-implantation. To assess the tissue response at this stage, slow and fast insertions were performed in animals that underwent recovery after surgery (n=3). Neuronal staining with Nissl and IgG labeling for BBB leakage was employed (**Fig. 5A**). We compared chronic fast insertions to acute fast insertions to analyze the temporal changes in BBB integrity over the course of a week.

**Figure 5:**
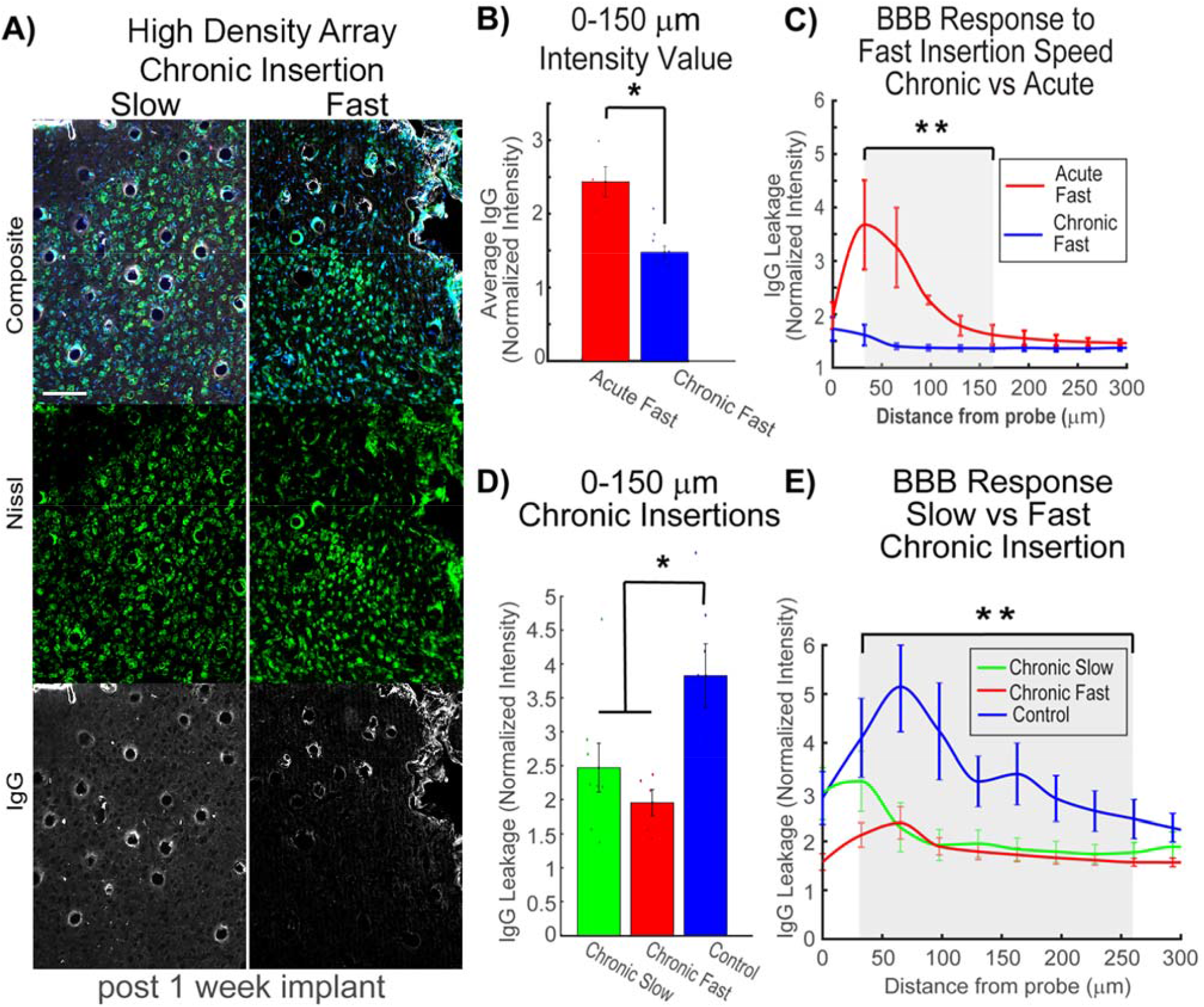
To test the chronic response of the High-Density Arrays has on the Blood-Brain Barrier(BBB), animals were implanted (N=3 rats/grp) for 1 week. **A)** Fluorescence staining for Hoechst stain (Blue); Nissl (Green); and IgG (White) displays the tissue response to the chronic insertion speeds (slow vs fast). The control speed was the acute slow insertion method. White scale bar = 100 μm. **B)** Normalized IgG intensity within 0 – 150 μm from probe hole for the acute (N=10) and chronic (N= 4) fast insertion speeds. **C)** Normalized IgG intensity within 10 μm bins up to 300 μm away from the probe hole for the acute and chronic fast insertion speeds. **D)** Normalized IgG intensity within 0 – 150 μm from probe hole for the chronic implants with varied insertion speeds (slow (N=8), fast (N=5), and control (N=7)). **E)** Normalized IgG intensity within 10 μm bins up to 300 μm away from the probe hole for the acute and chronic fast insertion speeds. ^*^ indicates p<0.05 and ^**^ indicates p<0.01 with an unequal variance t-test with Bonferroni correction for repeated measure.

Analysis revealed a significant disparity in IgG levels between acute and chronic conditions within 0 to 150 μm of the insertion site one week post-implantation (unequal variance t-test; p<0.05) (**Fig. 5B**). Overall normalized intensity values indicated a plateau in the BBB response at around 150-200 μm from the implant site. However, chronic fast insertions plateaued at approximately 60-100 μm from the implant site, with the highest levels observed near the baseline (200 μm away). Two-way ANOVA revealed significant effects of insertion group (p < 0.01), distance (p < 0.01), and the interaction between insertion group and distance (p < 0.01) on IgG leakage. Subsequent Tukey’s post-hoc tests demonstrated a significant increase in IgG for acute animals from 25-75 μm compared to all other groups (**Fig. 5C**).

Further investigations were conducted by implanting animals for one week using various insertion speeds (slow, fast, and control). The acute control involved the slow insertion Utah array. IgG intensity values were measured within 0-150 μm of the insertion site, indicating that both chronic slow and fast methods resulted in lower BBB leakage compared to the control after one week of implantation (p<0.05) (**Fig. 5D**). The overall results for normalized IgG intensity values suggested that the high-density arrays with both slow and fast insertion methods had a reduced impact on the BBB compared to the control Blackrock Arrays. Two-way ANOVA also identified significant effects of insertion group (p < 0.01), distance (p < 0.01), and the interaction between insertion group and distance (p < 0.01) on IgG leakage. Tukey’s post-hoc tests indicated a significant increase in IgG for acute animals 50-100 µm compared to all other groups (**Fig. 5E**). Taken together, the results from the chronic insertion experiments underscore the nuanced interplay between insertion speed and chronic BBB leakage, emphasizing the critical role of controlled insertion strategies in mitigating chronic BBB damage from high-density electrode implantation.

### Single wire insertions and recordings in Rats

Having demonstrated successful implantation of the FµA into the brain, we then investigated whether the electro-sharpening process affected the electrodes’ ability to record neuronal action potentials in layer 4 of the primary visual cortex of rats. Acute single wire recordings were conducted on 20 μm wires, confirming their capability to capture characteristic neural activity within the V1 cortex and respond to visual stimuli, as evidenced by robust neuronal single-unit waveforms (**Fig. 6A,B**) and responsiveness to drifting grating stimuli (**Fig. 6C**). Additionally, impedance values before and after the experiment remained stable and within the optimal range for acute neural recording settings (∼ 0.17 Mohm at 1 kHz).

**Figure 6:**
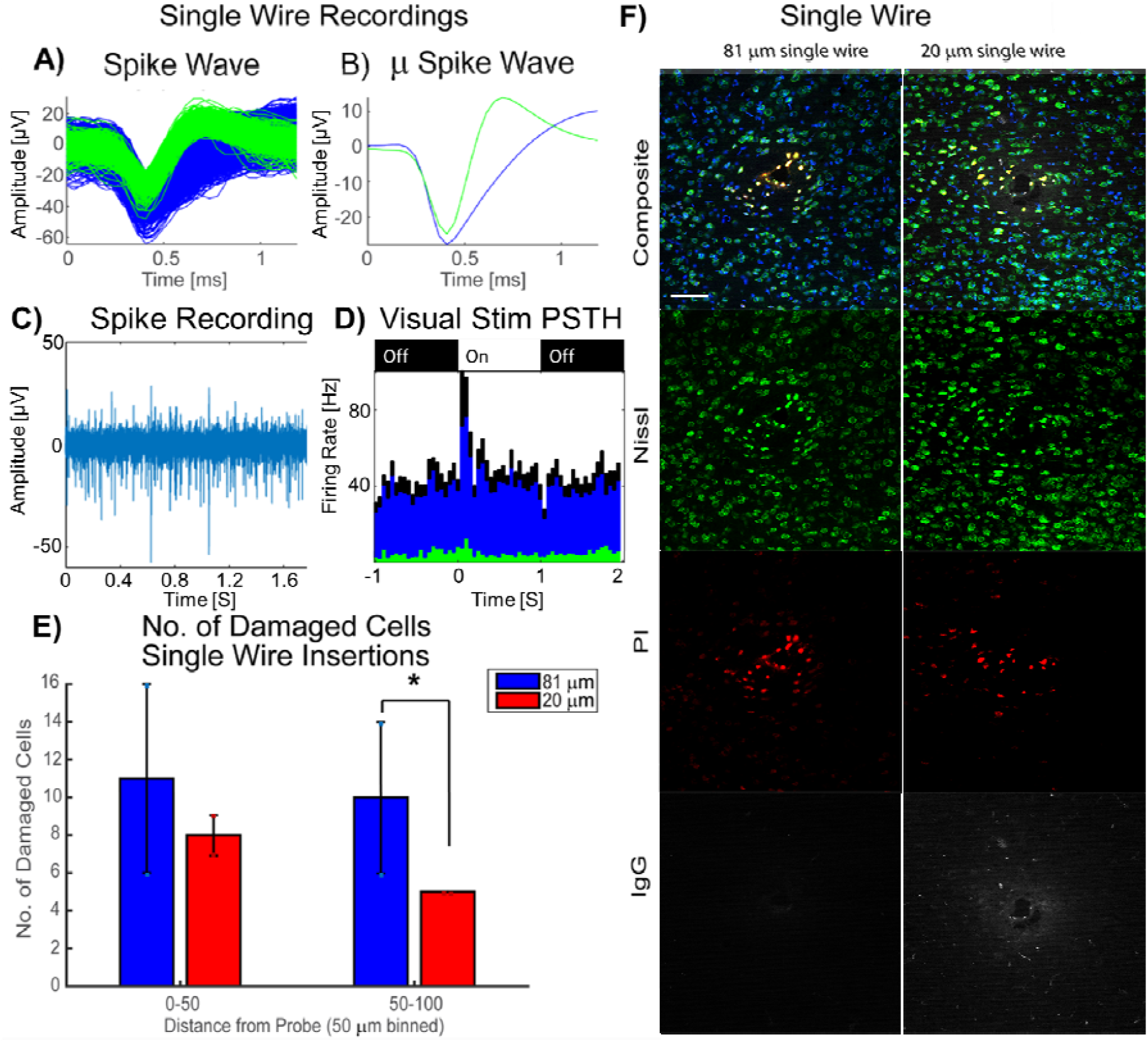
**A)** Single wire (20 µm) recordings from the HDA were inserted 520 μm down into the visual cortex (V1 area of the cortex). The electrode recorded cortical activity while a visual stimulus was played. The raw spike trace for an isolated neuron firing (green) and multi-unit activity (blue) was recorded. **B)** Mean Spike Wave form blue and green was measured. **C)** Raw spike stream filtered between 500 and 3000 Hz. **D)** Average peri-stimulus time histogram of 64 visual stimulation (On) to the contralateral eye showing evoked activity at stimulation onset at 0 s and offset response at 1 s. Green and Blue indicate spikes sorted from (A-B) and black indicates unsorted multiunits activity. **E)** The number of damaged cells (PI+) measured up to 100 μm away from the probe hole and averaged within 50 μm bins. **F**) Fluorescence staining for Hoechst stain (Blue); Nissl (Green); PI (Red); and IgG (White) displays the tissue response to the different single electrode diameter (81 μm vs 20 μm; N = 2 rats/electrode/bin). The yellow arrows identify areas of co-localization for the Nissl and PI stained cell and indicated areas of damaged cells or neurons around the insertion site. White scale bar = 100 μm.^*^ indicates p<0.05 with unequal variance t-test.

Comparative analysis of recording wires between the experimental 20 μm FµA single-wire and the control 81 μm Microprobe wire revealed that the larger diameter wire exhibited a greater number of damaged cells at distances of 50 to 100 μm from the insertion site (unequal variance t-test; p<0.05) (**Fig. 6D**). This suggests that the larger diameter wires cause greater compression related damage to distant cells, while nearby the electrode (0-50 μm) the damage is saturated and/or has large variability. Histological images, stained with Nissl, Hoechst, propidium iodide, and Immunoglobulin G, highlighted the areas of damaged cells or neurons (**Fig. 6E**), confirming the reduced cellular damage associated with the 20 μm FµA single-wire implantation. Taken together, these results emphasize the effective recording capabilities and reduced tissue impact of the electro-sharpened 20 μm FµA wires, positioning it as a promising candidate for precise and reliable acute neural recordings.

## Discussion

The manufacturing and performance of Paradromics’ arrays have been previously reported elsewhere^69^. Our primary objective was to evaluate a range of array design and insertion parameters to identify optimal settings for achieving improved bio-integration of high-channel-count microelectrode arrays, with a particular focus on bypassing the bed-of-nails issue. Functional performances of implants require careful consideration of design space parameters and the viability of the underlying tissue^2,23,24^. The study extensively assessed different tip profiles and insertion speeds, shedding light on the interplay between these variables and their impact on array insertability and adjacent tissue damage, highlighting the critical role of the blood-brain barrier (BBB) in assessing the success of these implantations.

Previous research has demonstrated a strong relationship between BBB leakage and recording performance^89,90^, leading us to assess the extent of damage through acute quantification of BBB leakage (IgG) and cell membrane rupture (PI). During the device insertion process, the probe could potentially tear through neurites or strain the surrounding tissue, resulting in the rupture of nearby cell membranes^57^. Although PI selectively labels cells with compromised membranes, it remains unclear whether these labeled cells are capable of recovery and long-term survival, emphasizing the need for additional studies to ascertain the fate of these PI-labeled neurons. Nonetheless, the presence of PI-negative, GCaMP6-positive cells near the electrodes suggests the acute presence of viable neurons surrounding the electrode tips.

In the *in vivo* two-photon experiments, we implanted multi-shank (∼24 channel) arrays into the cortex at a 30-degree angle due to the constraints posed by the skull, objective, probe, and tissue scattering. Although only the top shanks were observable, while the deeper shanks were obscured, all shanks successfully inserted into the brain. However, because the arrays were not slanted, when implanted at a 30-degree angle, the deeper shanks inserted first. While this method provided a rapid screening for optimal array tips and spacing, it does not fully represent the insertion mechanics associated with a large bed-of-needle arrays, even though it enabled us to visualize viable GCaMP active neurons during the insertions.

Consequently, perpendicular acute insertions were carried out with larger arrays. Despite the fact that blunt tipped arrays could insert at an angle because the shanks inserted through the dural and pial surface row by row^91^, blunt-tipped arrays cause a “bed-of-nails” effect resulting in brain tissue dimpling. Notably, the substantial strain on the tissue was evident in the two-photon data, revealing a large number of PI-labeled cells with blunt-tipped arrays when compared to angled or electro-sharpened tips. Electro-sharpened microwire bundles effectively alleviated the bed-of-needles effect compared to other tip profiles in this study, providing critical insights into the benefits of such designs in reducing tissue strain during insertion.

Although the production of electro-sharpened microwires entails greater overhead costs compared to blunt or angled microwires, the study’s findings underscore their superior performance in mitigating tissue strain during insertion.

Highlighting the significant role of neurotechnology companies, it is essential to emphasize the dual responsibility of ensuring the safety of implanted subjects ‘at all costs’ and sustaining the viability of the company. Ethical considerations arise due to the harm inflicted on patients left with obsolete implants when companies fail^92^. In this study, however, the results clearly indicate that electro-sharpened microwires are both necessary for inserting into the brain tissue to enable neural recordings as well as significantly reduce dimpling related tissue injury. Subsequent work by Paradromics has therefore prioritized electro-sharpened microwires in their arrays^69^.

Contrary to expectations, the angled tip caused the greatest number of cells to experience membrane damage, whereas the blunt tip caused greater BBB damage. Our data demonstrate that electro-sharpened microwire bundles alleviate the “bed-of-nails” effect more effectively than other tip profiles studied. The evaluation of insertion speeds aimed to facilitate the insertion of angle-tipped arrays and avoid the need for electro-sharpening the tips. Intriguingly, pneumatic insertion led to reduced IgG leakage but resulted in a significantly higher number of cells showing damage, as indicated by PI labeling.

These findings suggest that the BBB/vascular wall may exhibit greater structural stability compared to individual cells around the implant site with pneumatic insertion speeds. This aligns with previous observations that it takes several days for single-units to be detectable on pneumatically inserted Blackrock arrays in non-human primates^31^. Consequently, the study emphasizes that fast insertion speeds represent a compromise to minimize acute BBB and cell membrane rupture.

The process of electro-sharpening the microwire tip leads to a significantly thinner recording interface region due to tapering, which could change the surface area and geometry of the recording sites. However, our results demonstrate that individual electro-sharpened microwires can effectively record visually evoked single-unit activity with high fidelity in the V1 region of rats. Thus, the alteration in geometry not only facilitated easier and reliable implantation of high-density arrays but also effectively maintained single-unit signal fidelity. Our results confirm that electro-sharpened arrays with fast insertion speeds can successfully insert into the brain. Furthermore, the study underscores the critical role that insertion speeds play in determining the extent to which the BBB is compromised during implantation. Interestingly, the findings revealed no positive correlation between BBB leakage and initial cell membrane damage when varying insertion speed. However, it remains imperative to evaluate the long-term impacts of cell membrane damage. Nonetheless, the study presents promising parameters that can be leveraged in the development of a high-density, high-channel-count array. The evaluation of scalable electro-sharpened arrays with higher density in future studies is warranted.

### Limitations of this study

Several limitations are acknowledged in this study. The evaluation focused on the first 5 generations of prototypes of the Paradromics high-density arrays, resulting in a limited number of device replicates and incomplete exploration of the design space. Despite these constraints, the study yields statistically significant outcomes, offering insights into the impact of tip geometry, insertion speed, and microwire diameter on the cellular-level spatial scale of neuronal membrane integrity, BBB leakage, and successful cortical tissue insertion (Fig. 2-6). Future investigations should consider quantifying insertion force using ultra-sensitive sensors^47,50^, especially one that is compatible with a pneumatic inserter.

Another limitation associated with this study was its use of mice and rats. The choice of mice was essential for visualizing how electrode tip geometry impacts neuronal cell membrane injury under a two-photon microscope, providing novel insights into the spatial scale of cellular injury. However, the thinner dura and pia in mice, compared to clinically relevant thickness, result in less required insertion force for penetration, while the in vivo two-photon imaging necessitates a 30-degree angled insertion, increasing brain surface contact with array tips and requiring greater insertion force.

In contrast to mice, rat dura has a comparable thickness to human pia^93-96^. While the rat model efficiently excludes prototype designs that are unable to perpendicularly penetrate the brain, the assessment of higher channel count arrays necessitates evaluation in large animal models. Paradromics, building on insights gained from the mice and rats reported in this study, has advanced this research by developing a 30,000-channel array. Successfully implanted in a sheep model, this array demonstrated single-unit recordings on hundreds of channels^69^. Despite these advancements, the present study remains valuable for its contribution to fundamental basic science knowledge regarding the interplay between array design, insertion conditions, and the cellular spatial scale of tissue injury.

Finally, the rapid prototypes were insulated with relatively thick glass^70^. conventional hypothesis positing that high modulus materials like glass, ceramics, and carbon perpetuate micromotion in low modulus brain tissue, driving the foreign body response and glial scarring^97^, aligns with studies on flexible tethering to the skull contributing to glial activation^98,99^. However, extensive documentation highlighting the significant variability in glial scarring around stiff devices suggests the involvement of additional contributors to this undesirable tissue reaction^46,72,100-103^. For example, Rousche and Normann showed that two adjacent shanks with the same stiff material properties and insertion condition can generate highly disparate degree of foreign body response. Similarly, Williams et al showed that the same identical arrays can generate different tissue response and impedance spectra as well as recording performance^102,103^. Similarly, our prior work demonstrated large variability in tissue response and recording performance with identical silicon arrays^72,101,104^.

These results suggest there are other driving contributors^72,104,105^. Considerations should also extend to dielectric stability of low elastic modulus materials ^106^ and their contribution to performance as well as the fact that decreasing the cross-sectional area of implants can increase the overall flexibility of the device^37^. Similarly, it is crucial to note that not all glass and ceramics exhibit equal brittleness or flexibility^69,107^, and the materials used in the current Paradromics arrays differ from the rapid prototype devices used in this study^69^. Lastly, limited basic science knowledge exists on the collective impact of a large number of implanted shanks^69^ on long-term tissue response and device performance, necessitating careful evaluation due to the intricate interplay of design parameters^106^.

## Conclusion

This comprehensive study systematically evaluated various array parameters, including tip profiles and insertion speeds, to optimize the design of a high-channel-count microelectrode array for neural recordings. By carefully assessing the impact on tissue viability and the blood-brain barrier, we were able to identify key parameters that contribute to successful implantation and recording performance. The findings underscore the critical role of tip geometry and insertion speed in minimizing acute tissue damage, with electro-sharpened arrays and controlled insertion speeds demonstrating superior performance. These results lay a strong foundation for the development of future high-density arrays, promising significant advancements in both fundamental neuroscience research and the practical application of neural interface technology. Further investigations into long-term tissue responses and the scalability of these optimized parameters are essential to ensure the continued progress of this promising technology in the field of neuroprosthetics and cognitive neuroscience.

## Acknowledgments

This study is funded by the Defense Advanced Research Projects Agency’s (Arlington, VA, USA) Neural Engineering System Design program (DARPA-BAA-16-09-NESD-FP-001; Grant No. N66001-17-C-4005). The views expressed herein are those of the authors and do not represent the official policy or position of the Department of Veterans Affairs,

Department of Defense, or US Government. The authors would like to thank Paradromics Inc, especially Mina Hanna, Kunal Sahasrabuddhe, Robert J. Edgington, and Yifan Kong for providing devices, device images, and technical support for the V551-4B inserter, as well as X. Tracy Cui and Xu Pan for helpful discussions and help with the use of equipment.

