## Supplemental Data for "Electrode sharpness and insertion speed reduce tissue damage near high-density penetrating arrays"

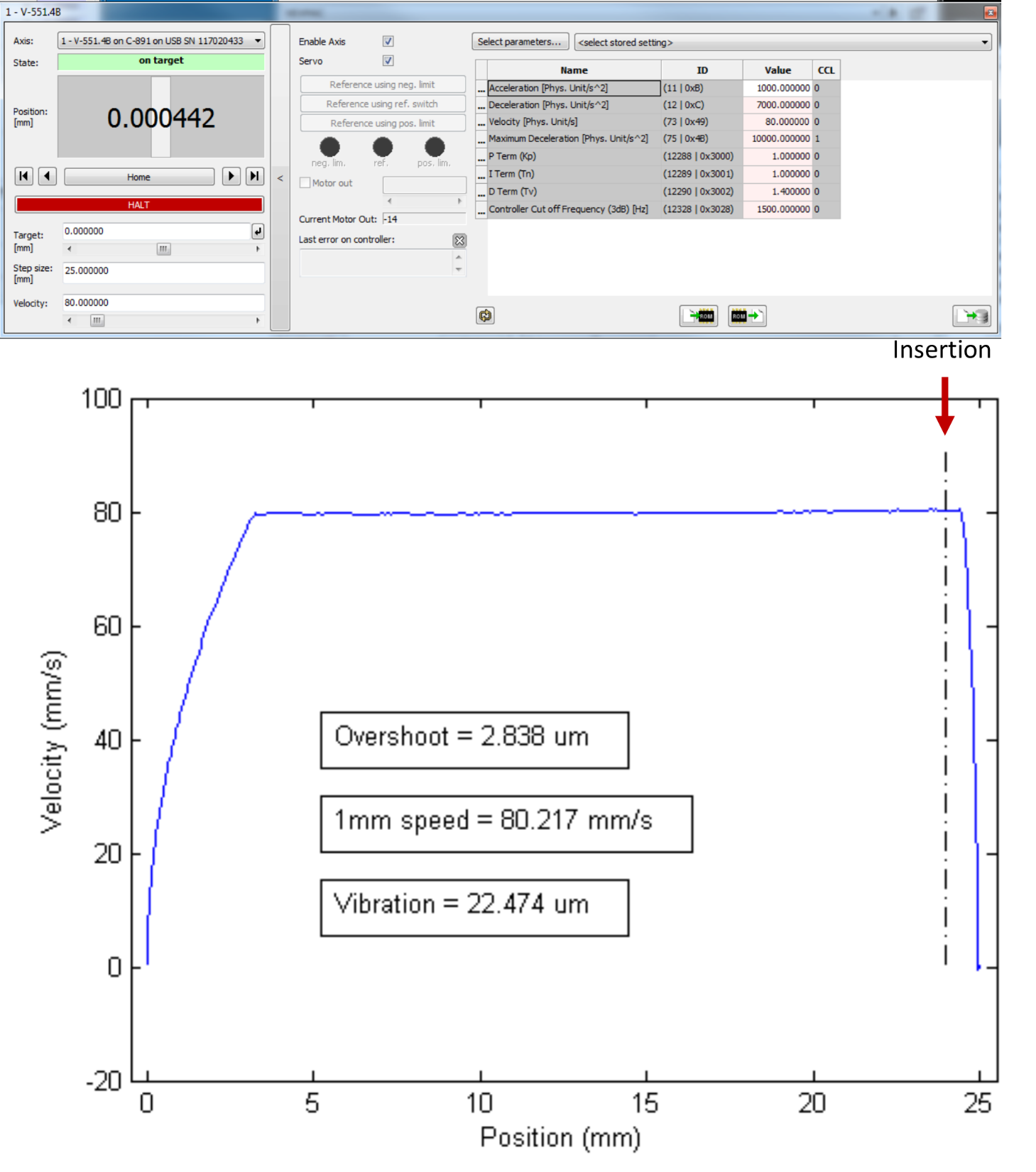
Supplemental Figure 1: V-551.4B calibration and performance. Top: C-891 controller inputs. Bottom: Velocity of the motor showing acceleration and deceleration profiles. Insertion into the brain occurs at the red arrow (-∙-∙ line).
